# BST2 induces vascular smooth muscle cell plasticity and phenotype switching during cancer progression

**DOI:** 10.1101/2024.09.10.612298

**Authors:** Caitlin F. Bell, Richard A. Baylis, Nicolas G. Lopez, Wei Feng Ma, Hua Gao, Fudi Wang, Sharika Bamezai, Changhao Fu, Yoko Kojima, Shaunak S. Adkar, Lingfeng Luo, Clint L. Miller, Nicholas J. Leeper

## Abstract

**Background:** Smooth muscle cell (SMC) plasticity and phenotypic switching play prominent roles in the pathogenesis of multiple diseases, but their role in tumorigenesis is unknown. We investigated whether and how SMC diversity and plasticity plays a role in tumor angiogenesis and the tumor microenvironment.

**Methods and Results:** We use SMC-specific lineage-tracing mouse models and single cell RNA sequencing to observe the phenotypic diversity of SMCs participating in tumor vascularization. We find that a significant proportion of SMCs adopt a phenotype traditionally associated with macrophage-like cells. These cells are transcriptionally similar to ‘resolution phase’ M2b macrophages, which have been described to have a role in inflammation resolution. Computationally predicted by the ligand-receptor algorithm CellChat, signaling from BST2 on the surface of tumor cells to PIRA2 on SMCs promote this phenotypic transition; in vitro SMC assays demonstrate upregulation of macrophage transcriptional programs, and increased proliferation, migration, and phagocytic ability when exposed to BST2. Knockdown of BST2 in the tumor significantly decreases the transition towards a macrophage-like phenotype, and cells that do transition have a comparatively higher inflammatory signal typically associated with anti-tumor effect.

**Conclusion:** As BST2 is known to be a poor prognostic marker in multiple cancers where it is associated with an M2 macrophage-skewed TME, these studies suggest that phenotypically switched SMCs may have a previously unidentified role in this immunosuppressive milieu. Further translational work is needed to understand how this phenotypic switch could influence the response to anti-cancer agents and if targeted inhibition of SMC plasticity would be therapeutically beneficial.

## Background

Through lineage tracing studies and epigenetic marker analyses, vascular smooth muscle cells (SMCs) have been found to possess profound phenotypic plasticity in response to disease stimuli.^1,2^ This has been best characterized in the pathogenesis of atherosclerosis, where in the setting of chronic, non-resolving inflammation, SMCs have been shown to lose their canonical marker gene expression (such as *Acta2* and *Myh11*), increase their proliferation and migration, and acquire unconventional protein expression that is more characteristic of alternative cell types.^1,3–5^ These diverse phenotypes appear to dictate the stability and pathogenesis of atherosclerotic plaques in both murine models and human disease.^3–7^ Additionally, one of the behaviors noted in several experimental contexts, though likely rare, is the ability of a SMC to acquire certain macrophage-like features.^8–10^

Less is known about SMC plasticity in the context of malignancy. Since Judah Folkman’s seminal work,^11–13^ investigators have sought to understand the cues that drive tumor vascularization and the mechanisms by which this phenomenon permits tumor growth.^14^ Vessel formation in solid tumors results from a multitude of microenvironmental signals between a wide range of cell types. Interestingly, there is significant vascular heterogeneity between tumor types, and even within the same tumor, which appears to have important implications for therapeutic responsiveness.^15–16^ Vascular SMCs have been shown to be a heterogeneous population that provide both pro-angiogenic signals and are critical for vascular stability and patency in the tumor.^17–19^ While changes in SMC marker expression have been reported,^20–22^ it is unknown whether the same degree of SMC phenotypic plasticity that exists within atherosclerotic lesions also is present within the TME, and, if so, whether this phenomenon could represent a new translational target for oncology.

## Materials and Methods

### Study Design

The prespecified objective of this study was to characterize SMC diversity and phenotypic plasticity during tumor angiogenesis. This was performed using several different indelible murine lineage-tracing models. Ex vivo mechanistic studies were performed using primary murine aortic smooth muscle cells. All experiments included at least three replicates with the actual “n” specified in each figure legend. Detailed methods are provided within the SI Appendix and are briefly summarized below.

### Mouse models

One SMC-lineage tracing model was developed specifically for this study, as detailed in SI Appendix and Supplementary Fig. 1A. These are tdTomato colored Myh11-CreERT2 ROSA-STOP-flox-eGFP lineage tracing mice. A single-color tdTomato reporter (Myh11-CreERT2 ROSA-STOP-flox-dtTomato) was also utilized. Only male mice inherit the *Myh11-Cre*^ERT2^ allele, thus any experiments performed using these mouse models were male mice that were heterozygous for the *Myh11-Cre*^ERT2^ allele and for the ROSA-STOP-flox-eGFP allele. The multi-color rainbow reporter mice (Myh11-CreERT2 ROSA-STOP-flox-Rainbow/+) have been used in prior publications.^5^ Myh11-Dre^ERT2^-Lgals3-Cre ApoE^-/-^ mice were provided by Dr. Gary Owens;^4^ this line was crossed with wildtype C57BL/6 mice from Jackson Laboratory to generate Myh11-Dre^ERT2^-Lgals3-Cre ApoE^wt/wt^ mice for these experiments. After treatment with tamoxifen, animals were rested for two weeks prior to tumor implantation as below. All animal studies were approved by the Stanford University Administrative Panel on Laboratory Animal Care (Protocol 27279).

### Cell lines

All cell lines were cultured according to the protocols provided by the suppliers. The C57BL/6 derived MC38, B16-F10, E0771, and LLC tumor cell lines were purchased from American Type Culture Collection (ATCC) and cultured in Dulbecco’s Modified Eagle medium (DMEM, Life Technologies), supplemented with 1% Penstrep and 10% FBS. Cells were maintained at 37C in 5% CO2 in a humidified incubator and passaged every 3-5 days.

### Subcutaneous syngeneic tumor models

2.5 x 10^5^ cells in 100ul volume of cold Hank’s Balanced Salt Solution were injected into subcutaneous flanks bilaterally. Tumor length and width was measured at days 5, 8, and 11 with calipers. Tumor volume = 0.5 x Length x Width^2^. Tumors were collected after 11 days of tumor growth and were weighed at time of collection.

### Tissue processing

Tissue for histologic analysis was placed in 4% paraformaldehyde, embedded in OCT, and sectioned at 10 um thickness. Tissue for FACS and/or scRNA-Seq was placed in cold PBS prior to tissue digestion. Tumor tissue was mechanically dissociated into 2-4mm pieces and then digested in 2.5 mg/mL Liberase and 0.1 mg/mL DNase (Roche) for 1 hour at 37C. Digested tissue was then passed through 70um filters and incubated with red blood cell lysis buffer for 10 minutes prior to neutralization with PBS. Cells were spun down in resuspended in PBS + 1% BSA for quantitative FACS or cell isolation prior to scRNA-seq. See SI Appendix, Methods for details regarding immunohistochemistry, microscopy, fluorescent image analysis, and FACS parameters.

### Single cell RNA sequencing

All single-cell capture and library preparation was performed at the Stanford Functional Genomics Facility and Stanford Genomic Sequencing Service Center. Cells were loaded into a 10x Genomics microfluidics chip and encapsulated with the barcoded oligo-dT-containing gel beads using the 10x Genomics Chromium controller according to the manufacturer’s instructions. Single-cell libraries were then constructed according to the manufacturer’s instructions (Illumina). Libraries were sequenced on an Illumina platform with targeted depth of 50,000 reads per cell for RNA. Fastq files from each library were aligned to the reference genome (mm10) individually using CellRanger Software (10x Genomics). Individual datasets were merged without subsampling normalization. The aggregated dataset was then analyzed using the R packages Seurat and CellChat, SI Appendix with further details. Lists of genes associated with each GO category were obtained from Geneontology.org, Panther analysis performed using the web-based platform.

### In Vitro Assays

Commercially available murine primary aortic smooth muscle cells (C57-6080; Cell Biologics) were grown on gelatin-coated cell culture surfaces (G1393: Sigma-Aldrich) in Dulbecco’s Modified Eagle medium (DMEM, Life Technologies), supplemented with 1% Penstrep and 10% FBS. Cell passages 3-6 used for in vitro experiments, confluence of 60-70% at initiation of protocols.

Cells cultured with murine recombinant BST2 (9940-BS-050; R&D Systems) at a final concentration of 0.5 ug/mL of protein for 48 hours unless otherwise stated. Except for proliferation assays, all assays performed in serum-deplete DMEM. TaqMan gene expression assays used TRIzol extraction, NanoDrop RNA quantification, and analysis on a QuantStudio 5 Real-Time PCR System. Proliferation assays and analysis performed in the Incucyte S3 Live-Cell Analysis System over a 4-day period. Cell-scratch assays used a P200 tip on gelatinized 12-well plates, analyzed using a validated Image J macro plugin.^23^ FACS-based phagocytosis assay as previously described^24^ using cultured murine SMCs as phagocytes and apoptotic MC38 tumor cells as target cells. Cells harvested for western blotting at predetermined time points pelleted and resuspended in Laemmli Sample Buffer (BioRad) and beta-mercaptoethanol, heated at 95C for 5 minutes. Membranes imaged on an Invitrogen iBright 1500 Imaging System, quantified on Image J. A full list of reagents, including antibodies, FACS parameters, and related experimental details provided in SI Appendix, Methods.

### Statistical analysis

Differentially expressed genes in the scRNA-Seq data were identified using a Wilcoxon rank-sum test, as implemented in the Seurat package version 5.0.1. Results were analyzed by GraphPad Prism 10 for statistical differences between experimental groups. For qPCR, fold-change values were analyzed using two-way ANOVA with Greenhouse-Geisser correction and Tukey’s post hoc comparisons. Western blotting for ERK phosphorylation analyzed using two-tailed ratio Student’s t test. Proliferation and wound healing assay results analyzed using two-tailed Wilcoxon-signed rank tests with Welch’s correction. Phagocytosis assay results analyzed using two-tailed Student’s t test. shControl and shBST2 tumor differences in volume, weight, and cellular composition were analyzed by two-tailed Wilcoxon-signed rank tests with Welch’s correction. P values < 0.05 were considered significant.

### Data availability

The raw and processed single-cell RNA sequencing datasets are made available on the Gene Expression Omnibus (GEO) database, accession code GSE262942.

## Results

### Lineage-traced smooth muscle cells demonstrate phenotypic switching in subcutaneously implanted tumors

We used the well-established Myh11-CreERT2 lineage tracing system^2^ on a tdTomato background and crossed it with ROSA-STOP-flox-eGFP mice, generating tdTomato colored Myh11-CreERT2 ROSA-STOP-flox-eGFP lineage tracing mice (referred to as ‘two-colored Myh11 lineage-tracing mice’). In adult two-colored Myh11 lineage-tracing mice, all native cells express tdTomato at baseline, and following intraperitoneal tamoxifen injection, any cell expressing Myh11 will lose tdTomato expression and constitutively express eGFP thereafter (Supplementary Fig. 1A, 1B). Given the specificity of Myh11 at tissue homeostasis for vascular SMCs,^2^ this permanently marks all SMCs and their progeny even if expression of this perivascular marker is eventually downregulated during pathology.^25^

To examine whether SMC phenotypic switching occurs during tumor angiogenesis, we subcutaneously injected the colon cancer cell line, MC38, into the bilateral flanks of syngeneic two-colored Myh11 lineage-tracing mice (Fig. 1A). On histologic examination, as expected, SMCs expressed eGFP while all other native cells expressed tdTomato and tumor cells remained colorless (Fig. 1B). We found a progressive increase in SMC investment from small peripheral vasculature in the tumor over a growth period of 11 days (Fig. 1C) as the tumors grew in volume (Supplementary Fig. 1C). Similar progressive growth and vascularization was seen in subcutaneously implanted melanoma B16-F10, breast cancer E0771, and lewis lung adenocarcinoma (LLC) tumor cell lines (Supplementary Fig. 1D, E). By high resolution fluorescent microscopy, we identified loss of the canonical SMC marker ACTA2 in eGFP+ lineage traced cells throughout histologic sections of the tumors (Fig. 1D), a process sometimes referred to as ‘de-differentiation’ in the vascular literature.^8,26,27^ On the other hand, we did note the appearance of eGFP+ cells away from appreciable vasculature within the TME (Fig. 1E, F), suggesting migration of SMC away from endothelial networks into the tumor interstitium.

**Figure 1.**
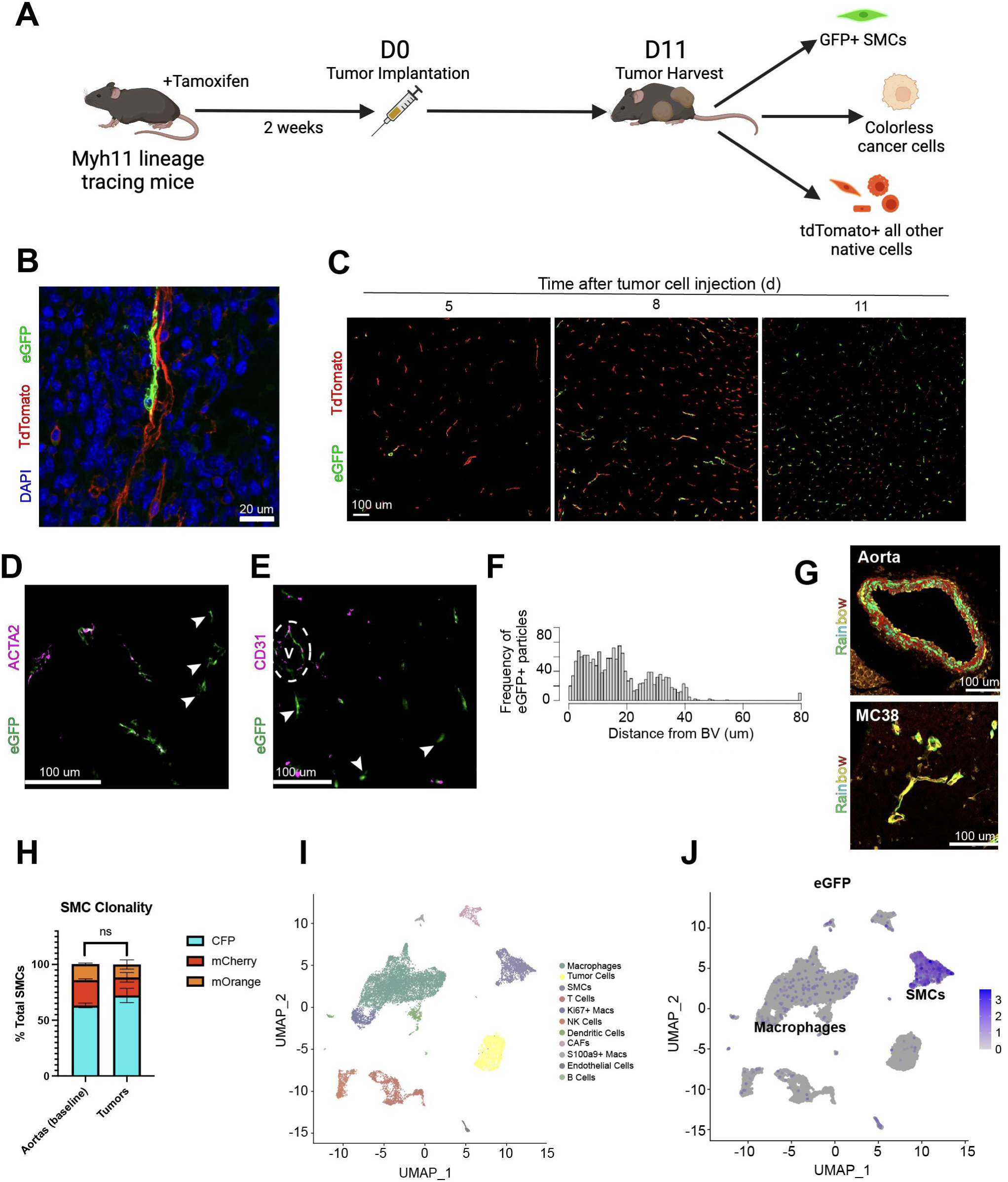
Phenotypic switching of lineage traced smooth muscle cells in subcutaneously implanted tumors. (A) Experimental design of tamoxifen administration to induce lineage tracing, rest, subcutaneous tumor implantation, and subsequent tissue collection. (B) Representative confocal microscopy image of MC38 tumor from two-colored Myh11 lineage-tracing mouse demonstrating native endothelial and immune cells (tdTomato) and an associated SMC (eGFP) in the environment of tumor cells (colorless). (C) Representative fluorescent microscopy images of MC38 tumors demonstrating progressive influx of native immune cells (tdTomato) and investment of SMCs (eGFP) in the TME over day 5, day 8, and day 11. (D) Representative fluorescent microscopy images of lineage-traced SMCs (eGFP) with and without coexpression of smooth muscle alpha-actin (ACTA2) after tumor harvest at 11 days. Arrows designate phenotypically switched SMCs. (E) Lineage-traced SMCs (eGFP) both in proximity with vascular endothelium (V) and without clear association (designated by arrows). (F) Computational analysis quantifying the proximity of eGFP+ particles to the nearest blood vessel (BV) in immunofluorescent images from subcutaneously implanted MC38 tumors on two-colored Myh11 lineage-tracing mice 11 days after implantation, n = 10 tumors. (G) Fluorescent microscopy images of aortic tissue and subcutaneously implanted MC38 tumor cell line from hemizygous multicolor Rainbow lineage-tracing mice. Central example vascular structure with pseudo-colored green endothelial cells and associated red SMCs (yellow), cyan-colored SMCs associated with endothelium in the upper right of the picture, scattered orange-colored SMCs seen throughout the histologic section. (H) FACS-based quantification of fluorescent SMC reporter frequency in aortic tissue versus implanted tumors from multicolor Rainbow lineage-tracing mice, n = 18 tumors. Not significant by two-tailed t test. (I) UMAP and unbiased clustering of scRNA-Seq data for all cells from 3 whole tumors and 10 tumors preferentially sorted for SMCs using FACS gating for eGFP+ cells, labeled with cell-specific cluster markers. (J) eGFP expression in the different cell clusters in (I). Fluorescent activated cell sorting = FACS. Single-cell RNA sequencing = scRNA-Seq. Smooth muscle cell = SMC. Tumor microenvironment = TME. Uniform manifold approximation = UMAP.

Dedifferentiation and subsequent clonal expansion of SMCs has been demonstrated in atherogenesis.^5^ To investigate whether these phenotypically switched and migratory cells were products of clonal expansion, we implanted the MC38 cell line into a hemizygous multicolor Rainbow SMC lineage tracer mouse model (Supplementary Fig. 1F).^5^ These mice have similar tamoxifen induced labeling of SMC but instead of uniform expression of eGFP, they stochastically acquire one of three fluorophores, such that dominance of one color over the baseline distribution is suggestive of clonality. Histologically, there were no clear areas around tumor vasculature with patches of clonal expansion (Fig. 1G), nor was there evidence of fluorescent marker expression skewing in tumor vasculature compared to baseline aortic smooth muscle cells (Fig. 1H).

To more precisely define the diversity of SMCs involved in tumor angiogenesis, scRNA-seq was performed on subcutaneously implanted MC38 tumors in two-colored Myh11 lineage-tracing mice resulting in four libraries; three libraries were biologic replicates from dissociated whole tumors, and one library used fluorescence-activated cell sorting (FACS) for eGFP+ cells from ten whole tumors to amplify the number of SMCs for analysis. We used a conservative cutoff of 6000 genes, as has been employed in other SMC lineage-tracing studies, to remove doublets and work to exclude events of cellular fusion or phagocytosis of one cell by another.^28^ Unbiased clustering and uniform manifold approximation projection (UMAP) of these datasets showed representation of all the expected cell types as identified by their gene expression profiles (Fig. 1I, Supplementary Fig. 1G, 1H). eGFP expression, as expected, was increased in the SMC cluster but also had notable levels of expression within the larger cluster of macrophages (Fig. 1J), representing 10% of eGFP cells in total.

Taken together, these data reveal evidence for SMC phenotypic switching during tumor angiogenesis, with cells losing contractile markers and readily migrating away from proper vascular structures. Initial studies using a Rainbow reporter suggest this process is not clonal in nature, and indicate that SMCs may upregulate genes typically associated with other cell types, as has been identified in other disease states.^1–10^

### Smooth muscle cells access transcriptional programs traditionally attributed to macrophages during tumor angiogenesis

To understand the diversity of SMC-derived cells present in this tumor model, all cells expressing eGFP transcript >=1 were subset and reanalyzed via principal component analysis and unbiased clustering, identifying eight distinct groups of lineage-traced cells (Fig. 2A). Each cluster has a distinct expression profile with analysis of differential gene expression (DGE) via Gene Ontology suggesting discrete biologic roles during tumor angiogenesis and interaction with the TME (Supplementary Fig. 2A). We then used the combined matrix with Monocle3 pseudotemporal analysis to attempt to map the trajectory of the SMC transition states (Fig. 2B). The smooth muscle trajectory appears to start with high contractile gene expression (eg, *Acta2*) which diminishes as SMCs take on a more proliferative and non-traditional phenotype. The cluster furthest from the original contractile cell state appears to take on a ‘macrophage-like’ state (based on marker gene expression), and upregulates pathways related to antigen-presentation and immune response relative to other lineage-traced SMC cellular groups (Fig. 2C, 2D, Supplementary Fig. 2B). Transcriptomic features of this cluster are not strictly aligned with an M1 or M2 phenotype, but rather are most like what has been previously described as a ‘resolution-phase’ macrophage (or type M2b) which is both enriched for antigen processing/presentation as well as immune cell clearance (Supplementary Fig. 2C).^29^ Additionally, expression of commonly utilized pericyte markers is upregulated in the larger cluster of cells on the UMAP representation, but not in the ‘macrophage-like’ cluster (Supplementary Fig. 2D).

**Figure 2.**
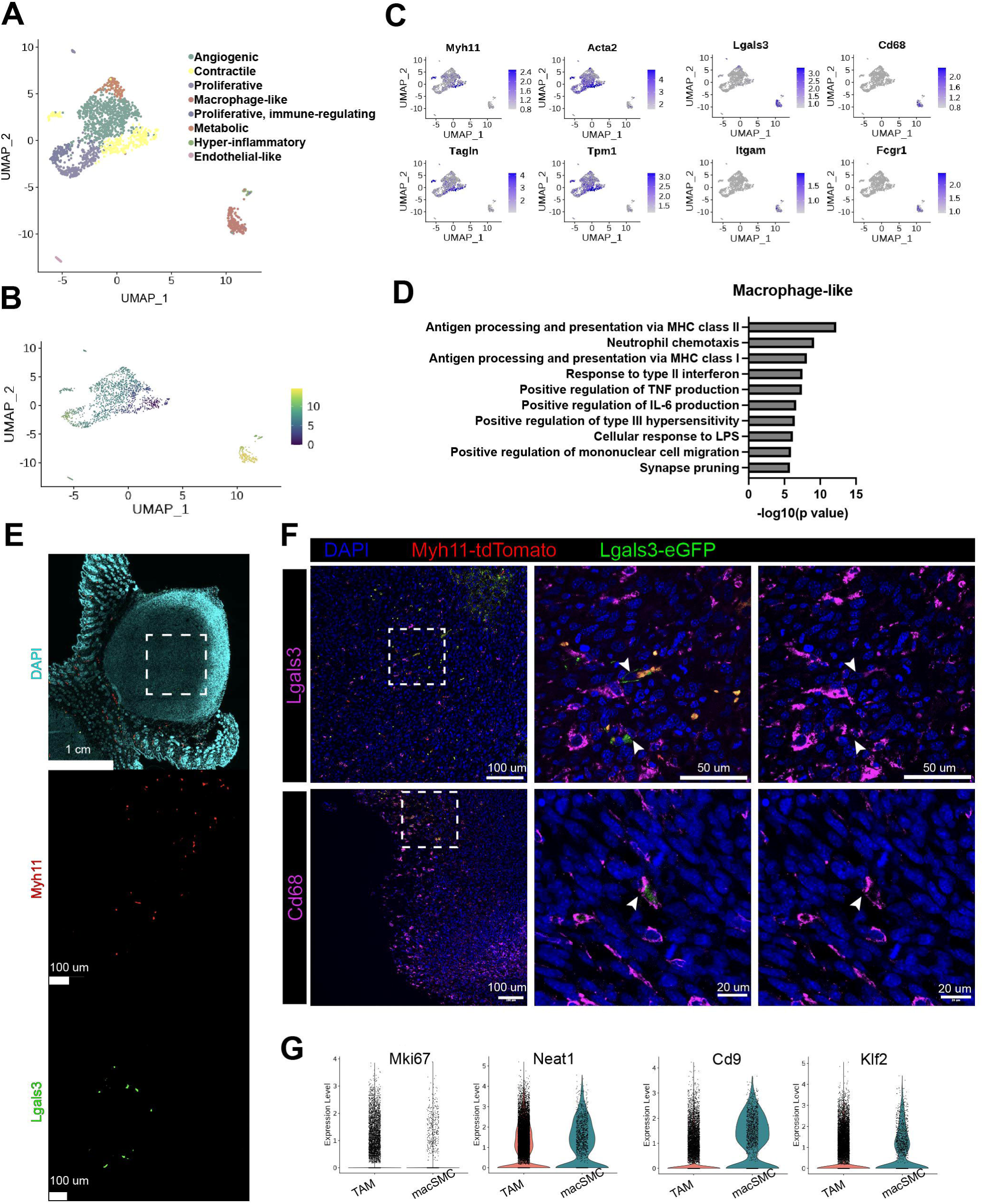
A subset of smooth muscle cells access transcriptional programs traditionally activated in macrophages during tumor angiogenesis. (A) UMAP and unbiased clustering of single-cell RNA sequencing (scRNA-Seq) data for all cells with eGFP >=1. Clusters annotated with functional names based on differential gene expression (DGE) analysis. (B) Pseudotime trajectory analysis of SMC scRNA-Seq data. (C) FeaturePlot of contractile gene expression (left) and macrophage gene expression (right) in UMAP of SMCs. (D) Biological processes Gene Ontology analysis of the macrophage-like cluster’s DGEs. (E) Representative fluorescent imaging of subcutaneously implanted MC38 tumors in dual lineage-tracing mouse model, and (F) confocal images of these tumors showing rare colocalization of LGALS3+ cells (green) with LGALS3 and CD68 staining (magenta). White arrows indicate rare cells with LGALS3+ lineage tracing and macrophage marker costaining. (G) ViolinPlot of transcripts with significant upregulation by non-parametric Wilcoxon rank sum test in SMC-derived macrophage-like cells compared to traditional tumor-associated macrophages. Differential Gene Expression = DGE. Single-cell RNA sequencing = scRNA-Seq. Smooth muscle cell = SMC. Uniform manifold approximation and projection = UMAP.

To confirm this phenotype-switching phenomenon, we next implanted tumors in a previously published dual lineage tracing mouse model that uses sequential activation of Dre and Cre recombinases to cause any Myh11+ SMC to change colors if they ever express *Lgals3*.^4^ Briefly, the Myh11 promoter drives a tamoxifen-inducible Dre recombinase, which removes a roxed stop cassette in the *Rosa* locus in front of a tdTomato-STOP-eGFP fluorescent reporter and a stop cassette on the *Lgals3* locus in front of an internal ribosomal entry site-Cre. If a cell subsequently expresses the macrophage-associated marker, *Lgals3*, it will express Cre, remove tdTomato, and express eGFP instead. Like the scRNA-seq data, eGFP+ cells (Myh11 expressing cells that subsequently expressed Lgals3) were found throughout the implanted tumors (Fig. 2E). However, like in atherosclerosis, only a rare subset of these SMCs colocalize with immunofluorescence staining for cell surface markers of true monocytes/stem cells/inflammatory cells perhaps suggesting that the Lgals3+ state is transient.^4^ There were rare instances of eGFP+ cells colocalizing with LGALS3 and CD68 suggesting that some of these cells can acquire the ability for protein level expression of monocyte/macrophage markers (Fig 2F).

We then directly compared the transcriptional signatures of traditional (non-SMC-derived) macrophages in the tumor to what will henceforth be referred to as ‘macrophage-like SMCs’ (macSMCs). The macSMCs had increased expression of *Mki67*, *Neat1*, *Cd9*, *Klf2, and Ezh2* among other transcripts (Fig. 2G). Neat1 has previously been implicated as a critical component in SMC phenotypic switching,^30^ Cd9 is a known marker of SMC proliferation in the atherosclerotic context,^31^ and Klf2 has a known role modulating vasculogenesis and blood vessel maturation.^32,33^ Ezh2 was more recently found to be an epigenetic repressor of contractile genes in phenotypically switched SMCs in aortic aneurysm models, beyond its traditional roles in PRC3 regulation.^34,35^ Pathway analysis of these DGEs in macSMCs highlight significant upregulation of cellular stress response pathways and cell cycle upregulation compared to traditional macrophages (Supplementary Fig. 2E).

These results demonstrate significant phenotypic plasticity and differential roles of SMCs investing in tumors after implantation. A subset of these cells loses detectable transcriptional signatures of SMCs and acquire markers thought to have roles in transitional states, including a macrophage-like cell. These macSMCs have a transcriptional signature suggesting more proliferation than their traditional macrophage counterparts,^36^ and retain transcriptional elements with established roles in vascular cell phenotypic switching and angiogenesis.

### BST2 expressed by tumor cells interacts with PIRA2 receptors on smooth muscle cells to activate macrophage transcriptional pathways and alter smooth muscle cell properties

To understand signaling inducing the phenotypic switch from a contractile SMC to a phenotype with certain features typically associated with macrophage-like cells, we utilized CellChat to predict ligand-receptor interactions related to this particular cellular subset (Fig. 3A).^37^ Comparing predicted ligand-receptor pairs between the tumor and typical tumor associated macrophages (TAMs) versus the tumor and macSMCs, revealed exactly one unique cell-cell interaction (Supplementary Fig. 3A, Fig. 3B): the cell surface receptor bone marrow stromal cell antigen 2 (BST2) on tumor cells binding to the cell surface receptor paired immunoglobulin-like receptor A2 (PIRA2) on macSMCs. This receptor interaction has previously been studied in dendritic cells as a negative feedback mechanism to type I interferon signaling, with PIRA2 expression predominating on macrophages and antigen-presenting cells.^38^ PIRA receptor complexes have been shown to activate macrophage pathways via phosphorylation of an immunoreceptor tyrosine-based activation motif, and then downstream via ERK, STAT1, and NFkB signaling pathways.^39,40^ BST2 is a type II transmembrane protein that is widely expressed and best known for its role in antiviral defense,^41^ but in more recent years has been found to be upregulated in several cancer subtypes and to promote tumorigenesis.^42^ Mechanistic work has implicated the receptor in promoting the ability of tumor cells to invade vasculature and metastasize.^43,44^

**Figure 3.**
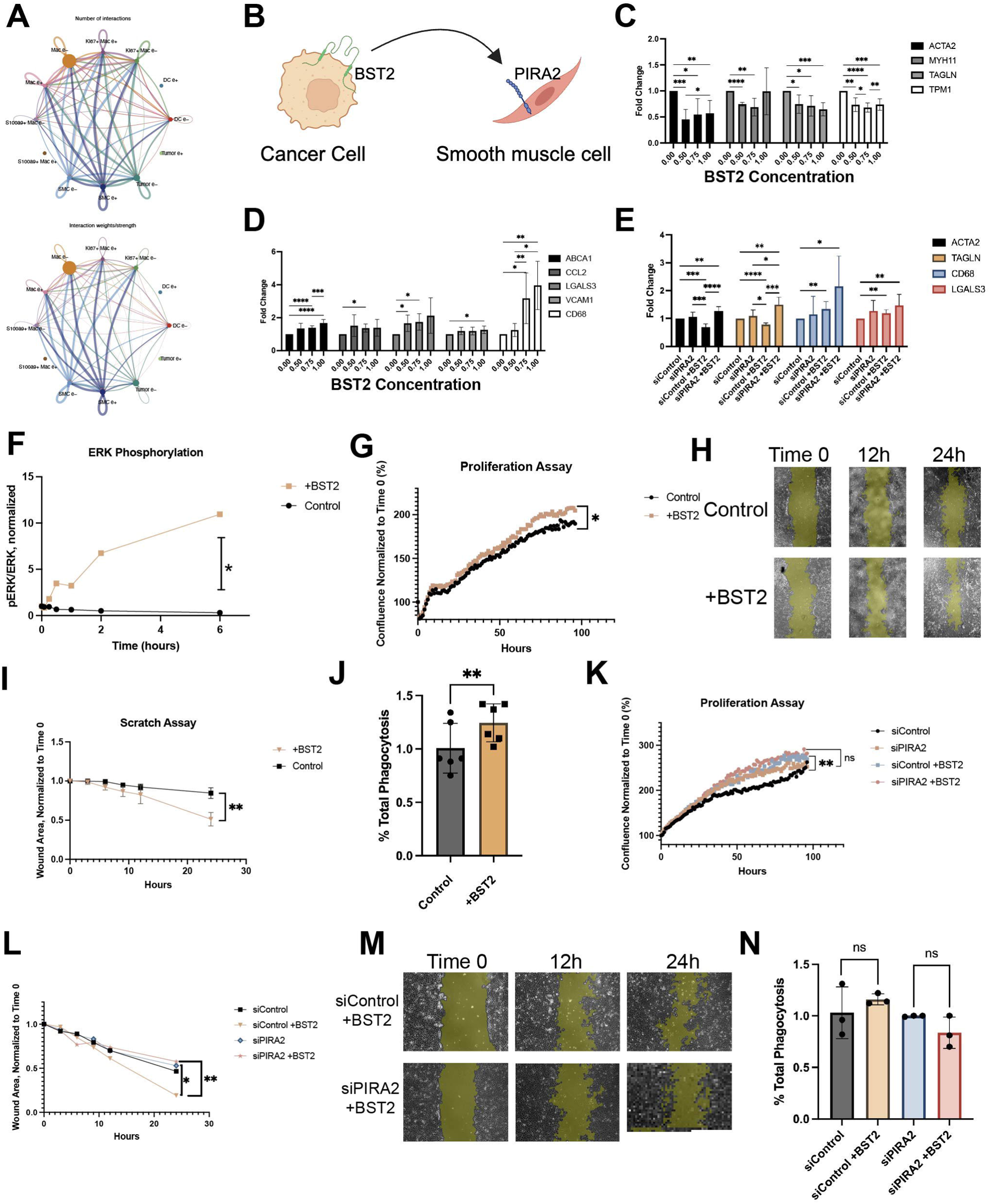
Macrophage transcriptional pathways on smooth muscle cells are activated by BST2 interacting with PIRA2. (A) CellChat was used to infer interaction partners between different cell clusters and GFP positive or negative transcriptional expression. netVisual_circle function used to display inferred numbers and weights of interactions between clusters of cells (B) Illustrative representation of the singular unique signaling pathway between tumor cells and SMC-derived macrophage-like cells versus tumor associated macrophages was via the BST2 cell surface receptor. (C) aSMCs were incubated with increasing concentrations of recombinant murine BST2 over 48 hours. RNA transcript expression for SMC markers and (D) macrophage markers was quantified by RT-qPCR and normalized to condition with 0.0 ug/mL BST2. n=9. (E) aSMCs transfected with either scramble or PIRA2-directed siRNA were incubated in the presence or absence of BST2, RNA transcript expression was quantified by RT-qPCR and normalized to the control condition without BST2. n=9. RT-qPCR results analyzed using two-way ANOVA with Greenhouse-Geisser correction and Tukey’s post hoc comparisons. (F) aSMCs were exposed to either BST2 or IgG, whole protein lysates were then harvested at prespecified time points for western blotting and probing of phosphorylated ERK (pERK) and ERK. Represented as the ratio of pERK to ERK over time, analyzed using two-tailed ratio Student’s t test. (G) Proliferation assay of cultured aSMCs in the presence or absence of BST2. Confluence normalized to condition at time 0, n=3, analyzed by two-tailed Wilcoxon-signed rank tests with Welch’s correction. (H and I) Wound healing assay of aSMCs in the presence or absence of BST2, wound area normalized to condition at time 0, n=4, analyzed by two-tailed Wilcoxon-signed rank tests with Welch’s correction. (J) FACS phagocytosis assay of aSMCs cultured in the presence or absence of BST2 and exposed to apoptotic MC38 cancer cells. % of total aSMCs phagocytosing cancer cells shown, n=6. (K) Proliferation assay of aSMCs transfected with either scramble or PIRA2-directed siRNA, in the presence or absence of BST2. Confluence normalized to condition at time 0, analyzed by two-tailed Wilcoxon-signed rank tests with Welch’s correction. (L and M) Wound healing assay of aSMCs transfected with either scramble or PIRA2-directed siRNA, in the presence or absence of BST2, n=4 per condition, analyzed by two-tailed Wilcoxon-signed rank tests with Welch’s correction. (N) FACS phagocytosis assay of aSMCs cultured in the presence or absence of BST2 and exposed to apoptotic MC38 cancer cells. % of total aSMCs phagocytosing cancer cells shown, n=3, analyzed by two-tailed Student’s t test. Bone marrow stromal cell antigen 2 = BST2. Fluorescent activated cell sorting = FACS. Murine aortic smooth muscle cells = aSMCs. Paired-Ig-like receptor A2 = PIRA2. Smooth muscle cells = SMCs. ns = not significant, *P<0.05, **P<0.005, ***P<0.0005, ****P<0.00005.

To test these in silico predictions, we cultured primary mouse aortic SMCs with escalating concentrations of murine recombinant BST2 and harvested the cells for RNA isolation. RT-qPCR found decreased expression of contractile genes and increased expression of macrophage-associated genes following exposure to BST2, consistent with the pattern observed in vivo (Fig. 3C, 3D). Co-culturing with recombinant BST2 was not, however, sufficient to induce significant changes in inflammatory markers or secretion of cytokines typically associated with an immune-activating TME (data not shown). Primary SMCs were then transfected with either a control or a PIRA2-directed siRNA (Supplementary Fig. 3B) and incubated either in the presence or absence of recombinant BST2. When PIRA2 was knocked down in SMCs, exposure to BST2 could not induce the same downregulation of contractile gene expression as in the presence of PIRA2 (Fig. 3E), though there was not a clear abrogation of effect with macrophage-associated genes. Time course analysis of primary murine SMC incubated in the presence of BST2 also showed an increase in ERK phosphorylation compared to control cells, consistent with its known activation cascade in immune cells that has been linked to macrophage phenotype switching^39,40^ (Fig. 3F, Supplementary Fig. 3C).

Taken together these experiments suggest that BST2 on the MC38 tumor cell may interact with PIRA2 on the SMC to promote a concurrent decrease in contractile markers, increase in traditionally macrophage-associated markers, and signaling cascades downstream of PIRA2 that have been associated with macrophage pathway activation. Consistent with its identification as a cell sharing certain features of the ‘resolution-phase’ macrophage, scRNA-seq data from the lineage-traced SMCs demonstrates a transcriptional signaling signature in the macSMCs more consistent with a mixed M1/M2 phenotype (Supplementary Fig. 3D).

To identify functional changes in the BST2-stimulated SMCs, further incubation assays were performed using primary aortic SMCs and recombinant protein. SMCs incubated with BST2 showed increased proliferation and migration by wound healing assay, which may relate to their penetration into the TME (Fig. 3G, H, I). When apoptotic MC38 cells were cocultured with SMCs, the presence of BST2 induced a modest but significant increase in the ability of SMCs to phagocytose apoptotic cancer cells by FACS phagocytosis assay (Fig. 3J), suggesting a possible functional shift toward nonprofessional phagocyte status. Using SMCs transfected with either control or PIRA2-directed siRNAs, the knockdown of PIRA2 eliminated both the proliferation and migration advantage seen in control SMCs (Fig. 3K, L, M). PIRA2 knockdown additionally reduced the ability of SMCs to phagocytose apoptotic tumor cells in the presence of BST2 (Fig. 3N). These data together suggest that signaling between BST2 and PIRA2 promotes SMC proliferation, migration, and enhancement of nonprofessional phagocytosis, along with loss of classic SMC contractile marker gene expression.

### Knockdown of BST2 in tumor cells decreases macSMC formation, smooth muscle cell investment within the TME, and tumor progression

To determine if this population of macSMCs is diminished in the absence of BST2, MC38 cells were treated with BST2-targeting shRNAs in a stable lentiviral transduction system to suppress BST2. A nontargeting shRNA (shControl) was used as well, demonstrating efficient suppression of BST2 expression (Fig. 4A). shBST2_1 was slightly more efficient than shBST2_2 and therefore used for all subsequent subcutaneous implantations. shBST2 and shControl-treated MC38 cancer cells were subcutaneously implanted in the bilateral flanks of syngeneic single-colored Myh11 lineage-tracing mice. Consistent with prior studies, ^43–47^ BST2-deficient tumors were smaller with reduced volume and weight at the time of harvest (Fig. 4B), and overall SMC content in the shBST2 tumors was significantly reduced (Fig. 4C, 4D). To account for differentials in tumor size, endothelial cell content of the tumor (as a surrogate of vascularization) was measured by CD31+ staining and cell segmentation and used as a corrective measure for SMC content; there were still significantly reduced SMCs in BST2-deficient tumors (Fig. 4E). In fact, there were so few SMCs in the TME of shBST2-treated tumors that it was difficult to quantify what seems to be a profound reduction in the ability of these cells to migrate into the tumors (Supplementary Fig. 4A). These data together suggest that tumor cell expression of BST2 is a key regulator in SMC tumor investment, even when correcting for differences in tumor size.

**Figure 4.**
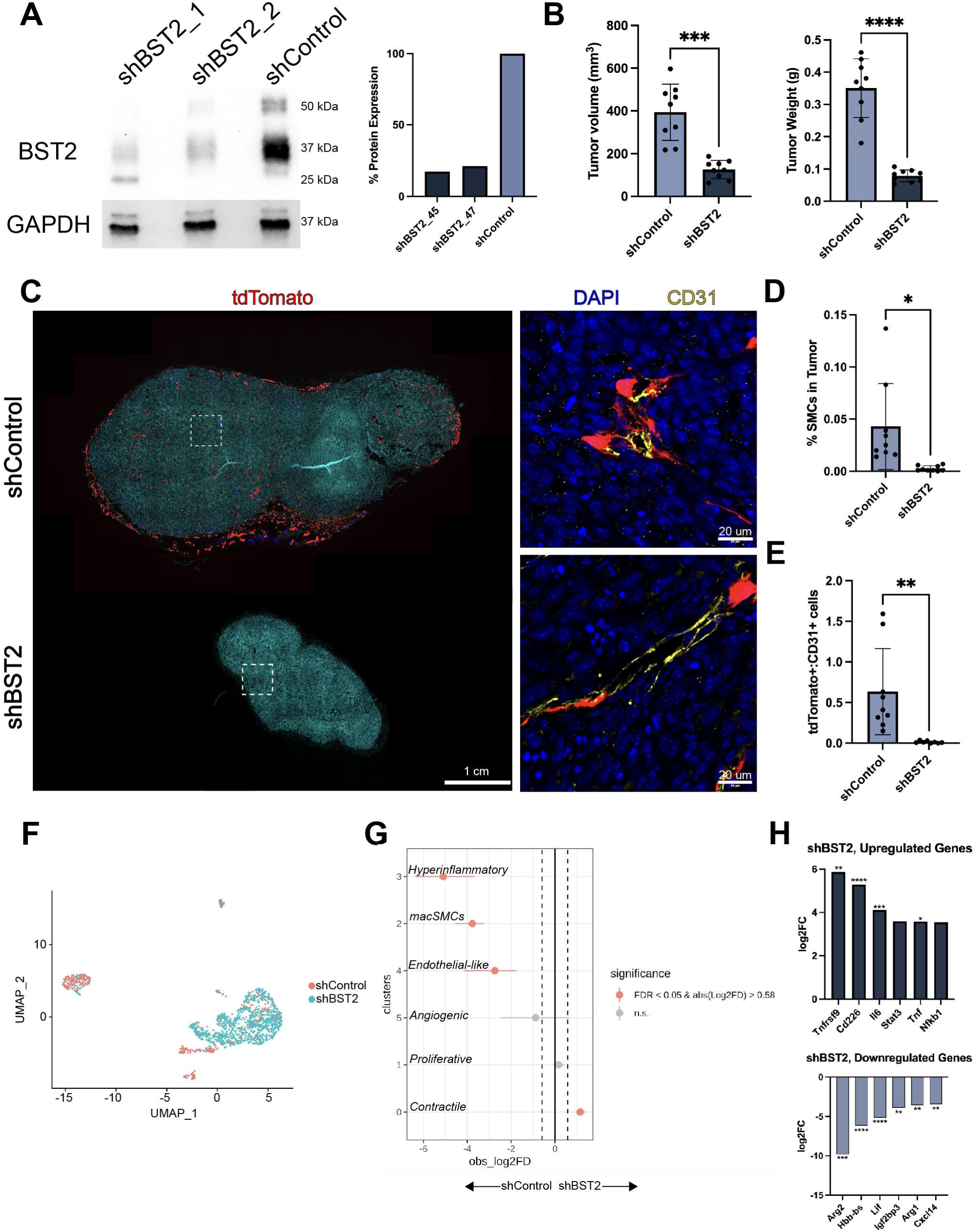
Knockdown of BST2 in tumor cells decreases smooth muscle cell tumor investment and phenotypic transition. (A) MC38 syngeneic colon cancer cell lines stably transfected with lentiviral vectors expressing short hairpin RNAs directed towards *Bst2* (shBST2) or a nonspecific control (shControl). Protein lysates probed for BST2 expression by western blot. (B) Tumor volume and weight of stably transfected tumor cell lines implanted into single color SMC lineage-tracing mice (Myh11-ERT-creT2 ROSA-STOP-flox-dtTomato), n=9, analyzed by two-tailed Wilcoxon-signed rank tests with Welch’s correction. (C) Representative fluorescent microscopy images of shControl and shBST2 MC38 tumors implanted in single color SMC lineage-tracing mice. IF for endothelial cells (CD31) performed with representative confocal microscopy images shown. (D) SMC frequency as a percentage of total cells in shControl and shBST2 tumors, n=9, and (E) ratio of tdTomato+ cells to CD31+ cells in shControl and shBST2 tumors, n=9. Analyzed by two-tailed Wilcoxon-signed rank tests with Welch’s correction. (F) UMAP and unbiased clustering of scRNA-Seq data for all cells with tdTomato >=1 from both shControl (blue) and shBST2 (pink) tumors. Clusters annotated with functional names based on DGE analysis. (G) Single cell proportion testing to compare the difference between the proportion of cells in clusters between shControl and shBST2 samples. Permutation testing used to calculate p values for each cluster. (H) DGE analysis between macSMCs from shControl and shBST2 tumors demonstrates M1 marker upregulation and M2 downregulation in shBST2 compared to shControl. Significant by non-parametric Wilcoxon rank sum test, expressed as log2(fold change) from baseline of 1.0 in control cells. Bone marrow stromal cell antigen 2 = BST2. Differential gene expression = DGE. Immunofluorescence = IF. Macrophage-like smooth muscle cells = macSMCs. Single-cell RNA sequencing = scRNA-Seq. Smooth muscle cell = SMC. Uniform manifold approximation = UMAP ns = not significant. *P<0.05, **P<0.005, ***P<0.0005, ****P<0.00005.

To understand how the absence of BST2 alters the phenotypic landscape of SMCs in the TME, tdTomato+ lineage-traced SMCs from shBST2 and shControl tumors underwent scRNA-seq, with unbiased clustering and UMAP projection (Supplementary Fig. 4B-D). This additionally provided an opportunity to use SMCs from a different fluorescent lineage tracing system to understand if the reporter system itself impacts our observations. Compared to SMCs in shControl-treated tumors, there were markedly fewer macSMCs present in BST2-deficient tumors and a higher proportion of SMCs in their contractile, quiescent states (Fig. 4F). This reduction in macSMCs and increase in classic, mature SMCs was statistically significant between the datasets (Fig. 4G). Direct comparison of macSMCs from control versus BST2-deficient tumors demonstrate that the presence of BST2 leads to a more immunosuppressive phenotype, as suggested by increased expression of Arg1/Arg2 and enhanced arginine metabolism (Fig. 4H, Supplementary Fig. 4E). The absence of BST2 resulted in a macSMC phenotype with upregulation of immune-activating factors and DGE pathway analysis suggesting regulation of SMC proliferation and vascular morphogenesis.

Assuming no additional unmeasured effect on traditional TAMs, these single cell analyses of in vivo lineage-traced SMCs in BST2-replete and deficient tumors suggest that BST2 enhances SMC plasticity and promotes a more immunosuppressive phenotype in the macSMC, which may contribute to BST2-dependent tumor growth.

## Discussion

Overall, these studies provide novel insights into the dynamic nature of SMC phenotypic switching during tumorigenesis. We find that, in addition to contractile and proliferative states, a small subset of SMCs can adopt a phenotype with some macrophage-like features in response to direct cell-cell interactions with the tumor interface. We observed that BST2 on the tumor cell, a known poor prognostic marker in both colon cancer and many other solid tumor subtypes,^45–47^ interacts with PIRA2 on the SMC to promote this phenotypic transition. This interaction provides an additional putative mechanism for how BST2 promotes tumor growth. BST2 has been shown to promote M2 phenotype skewing and an immunosuppressive TME.^48^ It is possible that the M2b-like signature of the macSMCs described in this study represent a previously unappreciated contributor to that immunosuppressive environment. With M2 macrophages largely regarded as pro-tumorigenic,^49^ the macSMC may represent a new translational target for those attempting to therapeutically shift the TME to an anti-tumor immunophenotype.^46^ While BST2 is known to directly enhance tumor cell proliferation, emerging data suggest additional roles in metastasis, invadopodia formation, and vascular invasion.^43,44^ These phenomena may be mediated in part by BST2’s newly recognized capacity to promote SMC migration away from developing neovessels and engraftment within the tumor interstitium, where they appear to participate in tumorigenesis.

Our findings provide novel insights into the role of SMCs in the TME, and further highlight the remarkable potential of SMCs to function beyond their classic contractile and vascular maintenance roles. Tumor angiogenesis has traditionally been viewed as pathological given its role in perfusion of the TME.^15,16^ However, our data suggest vascular cells may also directly contribute to tumor growth by undergoing phenotypic switching to cellular states that promote oncogenesis. While macSMCs only constitute a minority of the TAM compartment and a small percentage of tumor-associated vascular cells, their presence was seen in two separate lineage reporting models with scRNAseq analysis relying on transcriptional identification (rather than FACS alone) and conservative doublet thresholds. Evidence of their putative biological importance was provided by studies showing that suppression of the transition is associated with less aggressive tumor growth.

Notwithstanding these quality control measures, additional alternative hypotheses merit discussion. In addition to technical considerations related to microscopy, FACS sorting, and single cell analyses, it is possible that the observed cells which express genes found in both SMCs and macrophages may reflect fusion in the TME or represent macrophages which had recently phagocytosed an invading SMC. While we view these as less likely possibilities for reasons outlined above, these putative mechanisms cannot be definitively ruled out without lineage tracing experiments conducted with chimeric or parabiotic models.^50,51^

From a translational perspective, being able to redirect SMCs towards a more favorable phenotype could enhance tumor susceptibility to traditional chemotherapy, targeted treatment, or immunotherapy. There is prior work showing that phenotypic switching of pericytes, using pericyte-specific lineage tracing models, promotes tumor invasion and metastasis.^22^ Our study adds to the import of this phenotypic switching concept, highlighting an additional type of cellular change occurring. It also further suggests a role for targeting vascular compartment signatures, beyond what is possible with anti-VEGF agents. Such an approach is being pursued in atherosclerosis, where SMCs are being coaxed toward phenotypes that are cap stabilizing, anti-inflammatory, and reduce plaque vulnerability.^52^ The identification of a SMC that has some characteristics typically associated with macrophage-like cells builds upon this concept by finding a different type of phenotypic switching with characteristics suggestive of immunosuppression (and thus pro-tumor effect) that may represent a novel therapeutic target.

## List of abbreviations

BST2: Bone marrow stromal antigen 2
PIRA2: Paired Immunoglobulin-like Receptor A2
scRNAseq: Single cell RNA sequencing
SMC: Smooth muscle cell
TME: Tumor microenvironment

## Acknowledgments

C.F.B., R.A.B., and N.J.L. designed research; C.F.B., R.A.B., N.G.L., W.M., H.G., F.W., S.B., C.F., Y.K., S.S.A., L.L., and N.J.L., performed research; C.F.B., R.A.B., N.G.L., W.M., C.L.M, and N.J.L. contributed new reagents/analytic tools; C.F.B., R.A.B., N.G.L., W.M., H.G., F.W., and N.J.L. analyzed data; and C.F.B., R.A.B., and N.J.L., wrote the paper.

---

**Figure S1.**
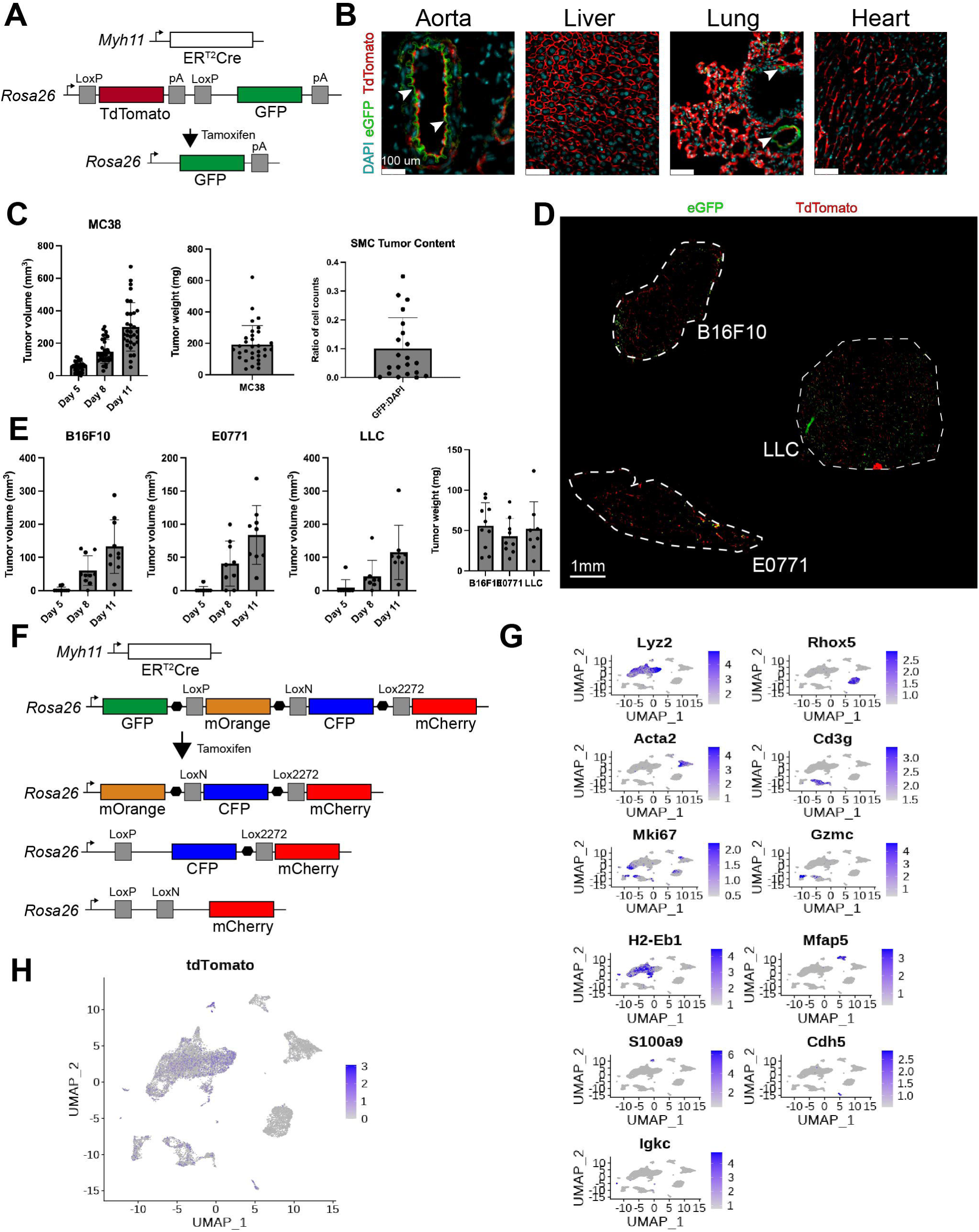

**Figure S2.**
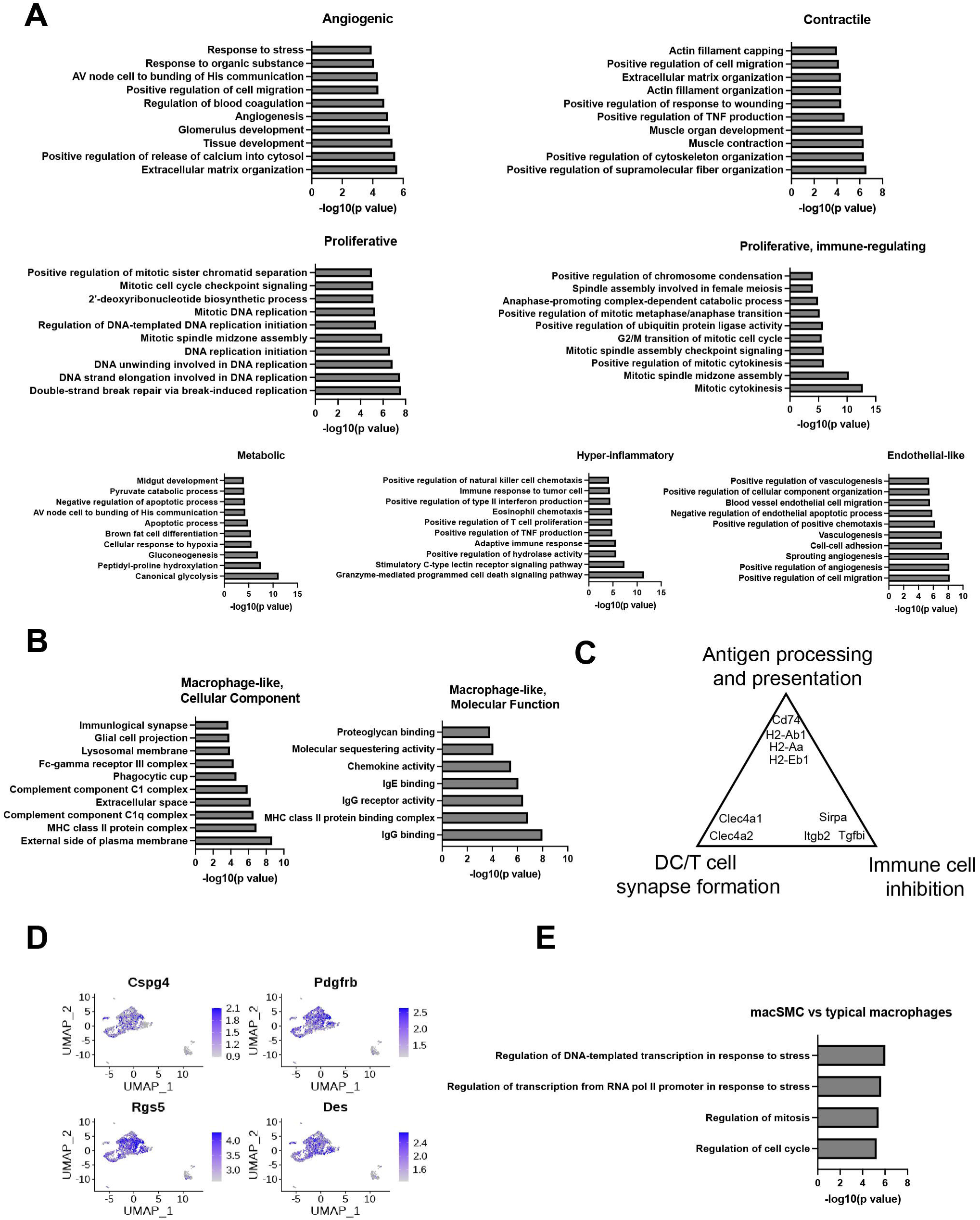

**Figure S3.**
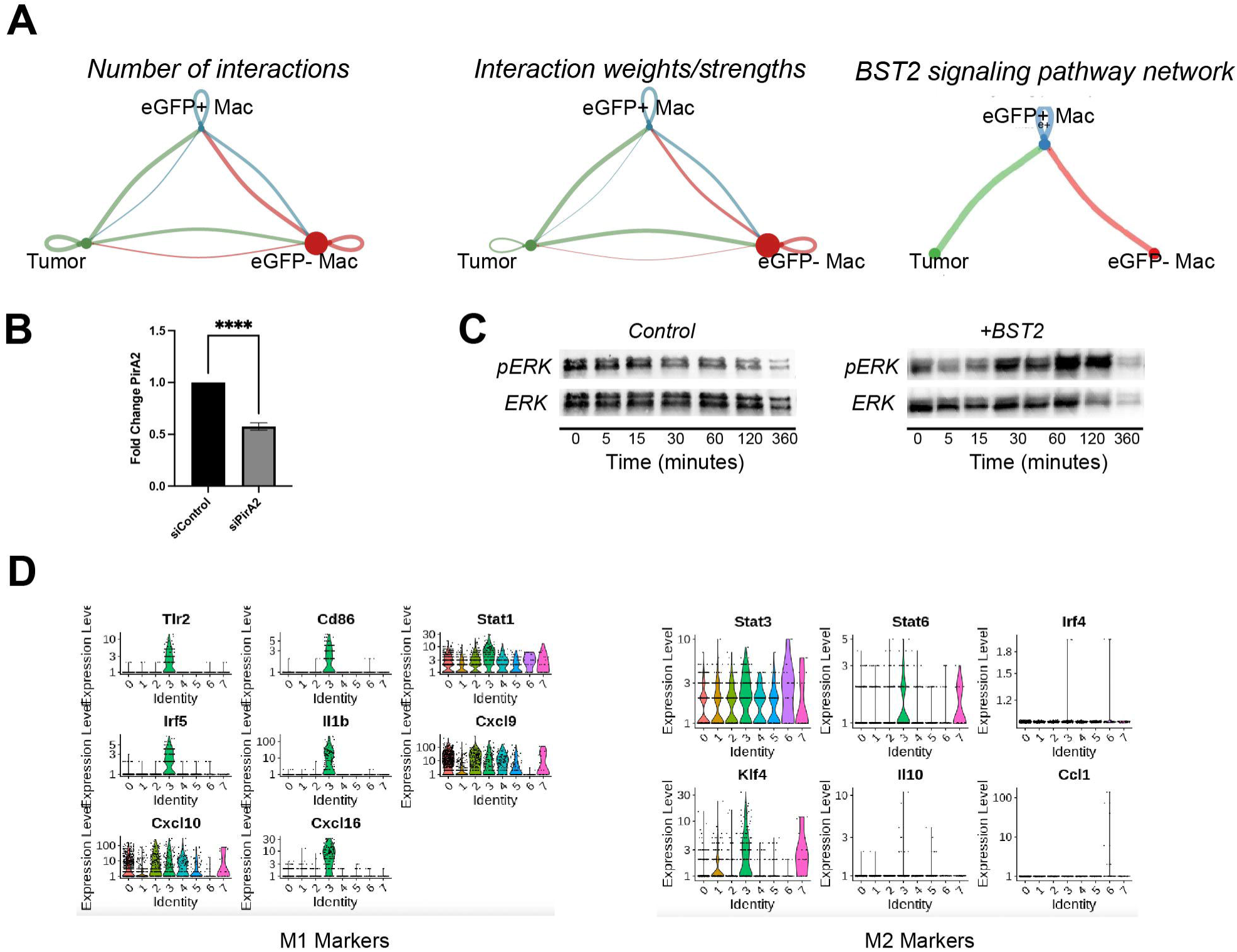

**Figure S4.**
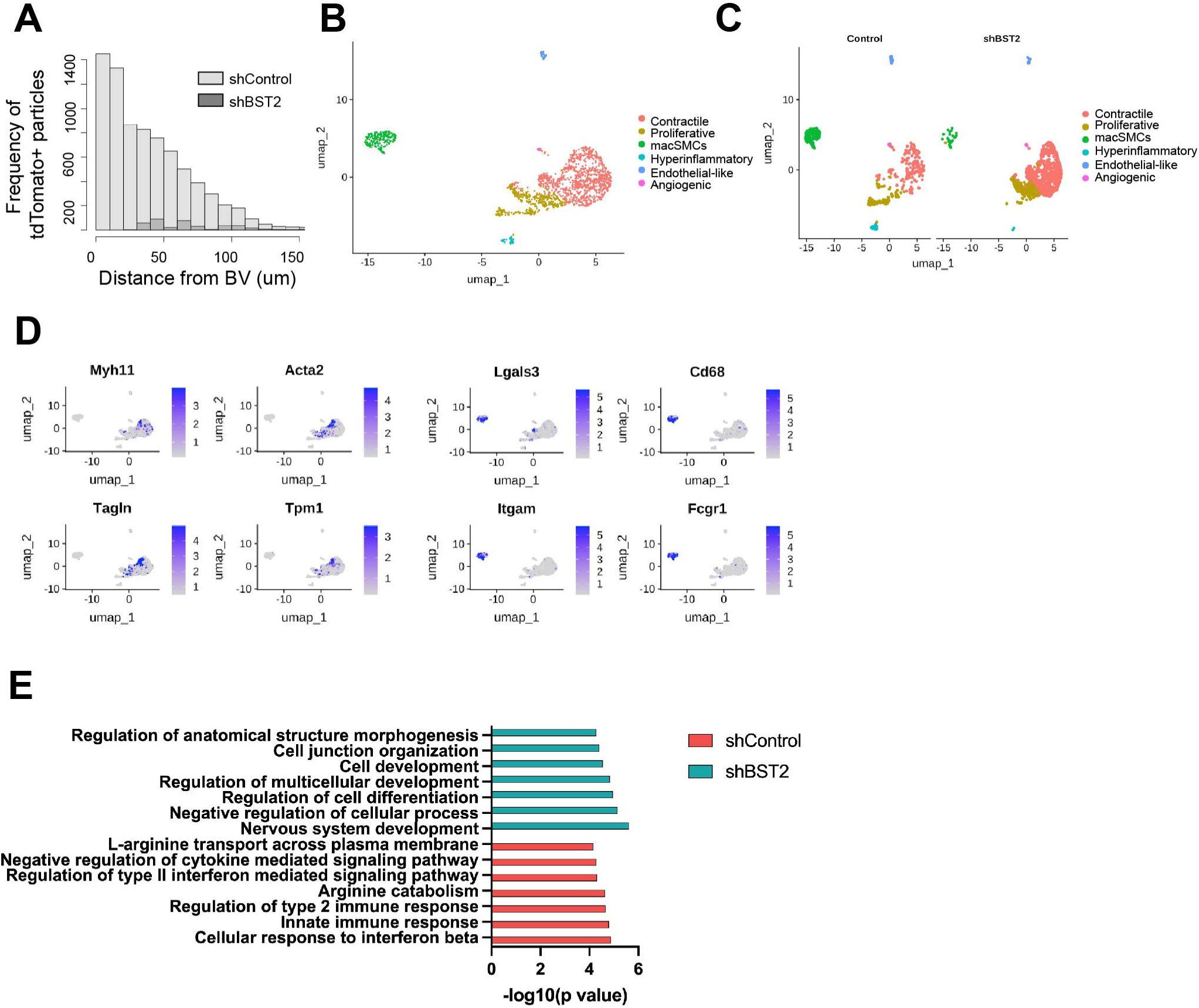

