## Supplemental Figure Legends for "BST2 induces vascular smooth muscle cell plasticity and phenotype switching during cancer progression"

**Supplemental Figure 1.** Phenotypic switching of lineage traced smooth muscle cells in subcutaneously implanted tumors. (A) Schema of the murine “two-colored” SMC lineage-tracing model. Before tamoxifen injection, all cells are labeled with tdTomato fluorescent protein. Tamoxifen injection induces Cre activity in the SMCs expression myosin heavy chain 11 (Myh11). Cre then cleaves LoxP sites flanking tdTomato and deletes the gene, allowing for expression of eGFP solely in mature SMCs while all other cells remain red. Transcription termination occurs downstream of the fluorescent transcript via 3’ polyadenylation sites (pA). (B) Representative fluorescent images of tissue sections in the two-colored mouse. eGFP restricted to SMC expression, indicated by arrows in the aortic media and pulmonary arterial tissue. tdTomato expression is seen diffusely in tissue types (vascular endothelium, hepatocytes, lung parenchyma, cardiomyocytes). (C) Tumor volume of subcutaneously implanted MC38 cancer cells over time, calculated as (tumor length x width x width)/2. Tumor weight measured at tissue harvest on Day 11. SMC tumor content measured histologically using computational cell segmentation and calculating number of eGFP+ cells to the total cell count in the histologic tumor section. (D) Representative fluorescent microscopy images of different types of syngeneic subcutaneously implanted tumors demonstrating native mouse cell (tdTomato) and SMC (eGFP) infiltration. Syngeneic lines are B16F10 (melanoma), LLC (Lewis lung adenocarcinoma), and E0771 (breast cancer). (E) Tumor volume of different subcutaneously implanted syngeneic tumor cells over time, and their weights at tissue harvest on Day 11. (F) Schema of the murine “Rainbow” multi-color SMC lineage-tracing model; hemizygous mice were used in all experiments. Before tamoxifen injection, all cells are labeled green with eGFP fluorescent protein (stop codon right after gene). Tamoxifen injection induces Cre activity in SMCs expressing Myh11. Cre then randomly selects flanking Lox cleavage sites, deleting gene fragments in between, allowing for stochastic expression of eCFP, mOrange, or mCherry; all non-SMCs remain green. The three colors will pass to the descendants of that cell, even if Myh11 is subsequently downregulated. Stop codons represented by black hexagons. (G) FeaturePlots showing gene expression of various markers used to inform cell identity of different unbiased clusters in the UMAP. (H) FeaturePlot of tdTomato expression in the different cell clusters in Fig. 1I. As expected, tdTomato expression is increased in clusters with likely immune cell identities and absent in the clusters of smooth muscle cells and tumor cells.

**Supplemental Figure 2.** A subset of smooth muscle cells access transcriptional programs traditionally activated in macrophages during tumor angiogenesis. GO analysis was used to gain insight into the biological significance of genes differentially expressed between SMC clusters. The top 100 genes for each cluster were utilized. See SI materials and methods for details, data available on GEO. (A) Biological processes GO analysis of DGEs from each additional lineage-traced smooth muscle cell (SMC) cluster. (B) Cellular and molecular GO pathway analyses of DGEs from the ‘macrophage-like’ cluster of cells. (C) Key features of resolution phase macrophages and their associated genes seen as significant DGEs in the macrophage-like cluster. (D) FeaturePlot of markers frequently used to look at pericytes. (E) Biological processes GO analysis of DGEs from SMC-derived macrophage-like cells compared to traditional tumor associated macrophages. Differential gene expression = DGE. Gene ontology = GO. Smooth muscle cell = SMC.

**Supplemental Figure 3.** BST2 interacts with PIRA2 to activate macrophage transcriptional pathways and alter smooth muscle cell properties. (A) CellChat receptor-ligand prediction algorithm results, demonstrating BST2 signaling from the tumor to GFP+ macrophages (i.e. macSMCs) but not to GFP- macrophages. Further methodologic details in the SI. (B) RT-qPCR of PIRA2 in smooth muscle cells (SMCs) transfected with a control siRNA or a PirA2-directed siRNA, n=12, ****P<0.00005, paired parametric two-tailed t test. (C) Western blot for ERK and phosphorylated ERK (pERK) SMCs in the absence or presence of recombinant BST2, cell lysates harvested at prespecified time points. (D) FeaturePlot of transcripts associated with M1 macrophage skewing and M2 macrophage skewing in single cell RNA sequencing of the GFP+ lineage-traced SMCs. The ‘macrophage-like’ cells are identity (aka cluster) 3 in green. Clusters 0 through 7 each correspond to smooth muscle cell clusters delineated in Figure 2A.

**Supplemental Figure 4**. Knockdown of BST2 in tumor cells decreases smooth muscle cell tumor investment and phenotypic transition. (A) Computational analysis quantifying the proximity of tdTomato+ particles to the nearest blood vessel (BV) in immunofluorescent images from orthotopically implanted MC38 tumors on single-colored Myh11 lineage-tracing mice 11 days after implantation, n = 9 tumors. (B) UMAP and unbiased clustering of single-cell RNA sequencing (scRNA-Seq) data for all tdTomato+ lineage-traced SMCs from both shControl and shBST2 tumors, labeled based on cluster-specific DGE. (C) UMAP separating shControl and shBST2 derived SMCs. (D) FeaturePlot of contractile gene expression (left) and macrophage gene expression (right) in UMAP of SMCs. (E) Biological processes Gene Ontology analysis of DGE between shControl and shBST2-derived macSMCs. Select significant genes from analysis in Figure 4H. Differential gene expression = DGE. Smooth muscle cell = SMC. Uniform manifold approximation = UMAP.
