## Supplemental Materials and Methods for "BST2 induces vascular smooth muscle cell plasticity and phenotype switching during cancer progression"

*Immunofluorescence*

Paraffin tissue sections were deparaffinized, rehydrated, antigen-retrieved (citrate), and blocked with fish skin gelatin PBS (6 g/L) containing 10% horse serum for 1 hour at room temperature. Slides were then incubated with primary antibodies shown in the below table overnight at 4C. Secondary staining with AF647 conjugated IgG was over 1 hour at room temperature before the slides were cover slipped.

| **Antibody** | **Catalog #** | **Dilution** |
| --- | --- | --- |
| DAPI | ThermoScientific, 62248 | 1:100 |
| ACTA2 | Abcam, ab5694 | 1:300 |
| CD31 | Abcam, ab124432 | 1:500 |
| LGALS3 | Cedarlane, CL8942AP | 1:500 |
| CD68 | Abcam, ab125212 | 1:500 |
| AF647 goat anti-rabbit IgG | Invitrogen, A21244 | 1:250 |
| AF647 goat anti-rat IgG | Invitrogen, A21247 | 1:250 |

*Microscopy*

For wide field microscopy, slides were imaged on a Leica Thunder DMi8. Lasers are 405/488/561/647, and objectives are HC PL FLUOTAR 20x/0.40 Dry and HC PL FLUOTAR 40x/0.6 Dry. For confocal microscopy, slides were imaged on a Ti2 AXR NSPARC Confocal. Lasers are 405/488/561/647, and objectives are 20x Plan Apo .80 Air, 60x Apo TIRF 1.49 Oil. Maximal intensity projection was used to generate representative images in the figures, Fiji was used to process and format the images.

*Fluorescent image analysis*

The original images were used for mean intensity measurements, and the background value was subtracted based on single-color and isotype control images. Entire tumor sections were analyzed, rather than selecting tissue ROIs. A computational pipeline was created for use with Fiji that optimizes cell segmentation by fluorescence given the often non-circular shape of tumor-associated vascular cells; circularity values for vascular cells were expanded to 0-1.00 to account for this in ImageJ. Images were uploaded for automated and uniform quantification of tumor area, total number of cells, number of lineage-traced smooth muscle cells, and number of cells staining for immunohistochemical marker of choice (e.g. CD31). The proximity of lineage-traced smooth muscle cells to blood vessels was computed by determining the radial distribution of CD31+ vascular structures to the fluorescent lineage traced cell (eGFP or tdTomato depending on the experimental model). Each marker positive lineage traced cell was analyzed to determine the closest CD31+ blood vessel, and this is the distance is reported in micrometers. Code can be found at https://github.com/jtmoore13/cell-dists.

*Fluorescent activated cell sorting (FACS) parameters*

Tissue processing prior to FACS discussed in main manuscript. All FACS analyses included gating on live, single cells with fluorescence-minus-1 controls. Dead cells were excluded by SYTOXTM Blue during sorting experiments (S34857, Thermo Fisher Scientific). A 100 um nozel was used.

Samples from (Myh11-ERT-creT2 ROSA-STOP-flox-Rainbow/+) multi-color rainbow reporter mice were analyzed on a Symphony (BD Biosciences) for quantification of eCFP, mCherry, and mOrange cells. Samples sorted for single cell sequencing (rather than whole tumor sequencing for some libraries) from Myh11-ERT-creT2 ROSA-STOP-flox-eGFP and Myh11-ERT-creT2 ROSA-STOP-flox-dtTomato lineage tracing mice were sorted on an Aria II instrument (BD Biosciences) for either eGFP or tdTomato depending on the smooth muscle cell lineage tracing line being used. Fluorescently sorted lineage-traced smooth muscle cells from 10 tumor cells during an experiment were then pooled for sequencing.

*Single-cell capture and library preparation and sequencing*

All single-cell capture and library preparation was performed at the Stanford Functional Genomics Facility and Stanford Genomic Sequencing Service Center. Cells were loaded into a 10x Genomics microfluidics chip and encapsulated with barcoded oligo-dT-containing gel beads using the 10x Genomics Chromium controller according to the manufacturer’s instructions. Single-cell libraries were then constructed according to the manufacturer’s instructions (Illumina), then sequenced on an Illumina platform with targeted depth of 50,000 reads per cell. Analysis of scRNA-Seq data Fastq files from each experimental time point and mouse genotype were aligned to the reference genome (mm10) individually using CellRanger Software (10x Genomics). Individual datasets were aggregated as needed using the CellRanger aggr command without subsampling normalization. The aggregated dataset was then analyzed using the R package Seurat version 5.0.1.

The dataset was trimmed of cells expressing fewer than 1000 genes. The number of genes, number of unique molecular identifiers and percentage of mitochondrial genes were examined to identify outliers. As an unusually high number of genes can result from a ‘doublet’ event, in which two different cell types are captured together with the same barcoded bead, cells with >6000 genes were discarded. Cells containing >15% mitochondrial genes were presumed to be of poor quality and were also discarded. The gene expression values then underwent library-size normalization and normalized using established Single-CellTransform function in Seurat. No additional batch correction was performed. Principal component analysis was used for dimensionality reduction, followed by clustering in principal component analysis space using a graph-based clustering approach via Louvain algorithm. UMAP was then used for two-dimensional visualization of the resulting clusters. Pseudotime trajectory analysis was performed using Monocle3 (version 1.0.0). Analysis, visualization and quantification of gene expression and generation of gene module scores were performed using Seurat’s built-in function such as “FeaturePlot”, “VlnPlot”, and “FindMarker.”

Differentially expressed genes identified in Seurat by the “FindMarker” function were used for pathway analysis. The top 100 differentially expressed genes in each cluster in order of fold change were used; Bonferroni corrected p values < 0.05 were considered significant, non-significant genes were not utilized. This gene list was input for pathway analysis be GO enrichment (PANTHER Overrepresentation test, version 18.0, Geneontology.org). FDR P <0.05 by Fisher’s Exact testing was considered significant. Top main hierarchical pathways reported in order of most significant p value.

Ligand-receptor interaction analysis was performed using CellChat^1^ (version 1.4.0). Seurat clusters from the original analysis were broken into eGFP transcript >= 1 positive and negative objects (annotated as e+ or e-). The processed Seurat object was fed into the ‘creatcellchat()’ function and processed using its standard pipeline. Briefly, the Seurat count matrices were extracted along with the Seurat/TS annotation generated previously. CellChatDB.mouse was loaded and differentially expressed genes and interactions were identified in the CellChat object via identifyOverExpressedGenes() and identifyOverExpressedInteractions(), respectively. The CellChat algorithm was then run to calculate the probably interactions and pathways via computeCommunProbPathway(). We also ran filterCommunication() to filter out interactions with less than 10 cells in each cell type. BioRender used to illustrate certain results for clarity.

*TaqMan Assays*

RNA was isolated from cell lysates using Trizol (Invitrogen) and quantified using a Nanodrop (Agilent Technologies). cDNA was generated using High Capacity cDNA Reverse Transcription Kit (4368814, ThermoFisher Scientific). Differences in gene expression across cDNA samples were examined and quantified using TaqMan gene expression assay and master mix (ThermoFisher Scientific). Amplification results were normalized to GAPDH internal controls and expressed as fold change in gene expression relative to the experimental control condition. A list of probes used is provided below. QuantStudio 5 Real-Time PCR System (ThermoFisher Scientific) used for generation of CT curves and analysis.

| **Assay Target** | **Assay ID** |
| --- | --- |
| ACTA2 | Mm00725412_s1 |
| MYH11 | Mm00443013_m1 |
| TAGLN | Mm00441661_g1 |
| TPM1 | Mm00445895_g1 |
| ABCA1 | Mm00442646_m1 |
| CCL2 | Mm00441242_m1 |
| LGALS3 | Mm00802901_m1 |
| VCAM1 | Mm01320970_m1 |
| CD68 | Mm03047343_m1 |
| PIRA2 | Mm02768273_g1 |

**In vitro assays:**

Key experimental materials in the below in vitro assays include:

- Primary murine aortic smooth muscle cells (Cell Biologics, C57-6080)
- Recombinant murine BST2 (R&D Systems; 9940-BS-050)
- siControl (Invitrogen, Silencer Negative Control, Assay #4404021)
- siPIRA2 (Invitrogen, Assay #284011)
- MC38 murine colon adenocarcinoma cell line (Kerafast)

*Smooth muscle cell transfection*

Cells were seeded on gel coated 6 well plates at 50-70% confluence (7500 cells per well) the night before transfection. Manufacturer’s instructions for use of Lipofectamine 3000 (Invitrogen) in transfection of siRNA was followed for siControl and siPIRA2 constructs. Cells were incubated for 48 hours before proceeding with analysis via the assays described below.

*Proliferation Assays*

Smooth muscle cells were seeded on gel coated 6 well plates at 50-70% confluence (7500 cells per well) the night before start of the experiment. Immediately before placing plates in Incucyte Live-Cell Analysis System (Sartorius), wells were rinsed with PBS and fresh growth medium was added. Recombinant murine BST2 was added to half of the experimental wells. The cell placte was placed into the Incucyte for 30 minutes prior to initial scanning. Settings used 10X objective, phase contrast, 16 point scan, with a scan interval of every 1 hour. Incucyte software was used to calculate density of area occupied by cells over time; density for each experimental well normalized to its own density at time 0.

*Scratch Assays*

Cell migration was assessed performing a scratch wound assay as previously described.^2^ Briefly, primary murine aortic smooth muscle cells were plated in a 6 well dish at 90% confluency and incubated overnight. Cells were then serum starved in DMEM + 3% FBS for 24 hours. Media was aspirated and a scratch wound was made across the center of each well using a P200 pipet tip. Cells were then incubated with medium either in the presence or absence of BST2. Differences in cell migration were captured using a phase contrast inverted microscope (Nikon). Scratch area at different time points was calculated using a previously published ImageJ plugin developed for high throughput analysis of scratch assays.^3^

*Phagocytosis Assays*

These assays examined phagocytic capabilities of smooth muscle cells, in the presence or absence of BST2, for eating apoptotic tumor cells. Primary murine aortic smooth muscle cells. Smooth muscle cells were plated in P10 dishes and pre-labeled with CellTracker Deep Red (C34565, ThermoFisher); then they were either incubated with or without recombinant BST2 for a full 24 hours. Syngeneic colon cancer cell line MC38 was plated in P10 dishes and was pre-labeled with CellTracker Green CMFDA (C2925, ThermoFisher). To create apoptotic labeled cells, they were incubated with 1um STS overnight.

Cells were harvested and counted. A ratio of 1:3 smooth muscle cells to tumor cells were plated in a 12 well plate and incubated for 2 hours before collection for FACS analysis. Co-expression of CellTracker Green and Deep Red signal was considered to be indicative of phagocytosis by the smooth muscle cells. Expressed as a percentage of the total labeled smooth muscle cells participating in phagocytosis (as known nonprofessional phagocytes).

*Western Blotting*

Samples of pelleted cells from culture prepared in Laemmli Sample Buffer (BioRad) + BME, heated to 95C for 5 minutes. 15 well Mini-PROTEAN TGX gel (Biorad) loaded with Precision Plus Protein Kaleidoscope (Biorad) and 10 ul of each prepared sample, run in Tris/glycine/SDS buffer at 100V for 1 hour. Gel transferred to PVDF membrane (Biorad). Membrane blocked with 5% BSA in 1x TBST for 1 hour before applying primary antibody to the membrane overnight at 4 degrees. After washing with 1x TBST, secondary antibody applied at room temperature for 1 hour, and then subsequently washed again. Membrane developed with SuperSignal West Pico PLUS Chemiluminescent Substrate (ThermoFisher, 34579) and imaged on an Invitrogen iBright 1500 Imaging System, quantified on Image J.

| **Antibody** | **Catalog #** | **Dilution** |
| --- | --- | --- |
| BST2 | Abcam ab246508 | 1:1000 |
| GAPDH | Santa Cruz sc-32233 | 1:1000 |
| ERK 1/2 | Abcam ab184699 | 1:1000 |
| phospho-ERK 1/2 | Abcam ab278538 | 1:1000 |
| Anti-rabbit IgG, HRP-linked | Cell Signaling Tech, #7074 | 1:4000 |

*Generation of BST2-deficient MC38 Cell Lines*

1.6 x10^4^ MC38 cells were seeded per well in a 96 well plate, triplicate wells per lentiviral construct. The following day the cells were incubated with 8 ug/mL of hexadimethrine bromide (Sigma Aldrich), and 10 uL of lentiviral particles were added to the appropriate well for overnight incubation. The control vector was pLKO.1-puro-CMV with a non-targeted shRNA control (Sigma-Aldrich SHC016), and the BST2-targeting vectors were in the same backbone with short hairpin construct directed against BST2 (Sigma-Aldrich clone IDs 125455 (aka BST2_1) and 125457 (aka BST2_2)). Cells were given fresh medium without puromycin the following day, then medium was replaced with DMEM +FBS +Pen/Strep + 4 ug/mL of puromycin as was predetermined by a puromycin kill curve in this cell line. Medium was replaced every 3-4 days, cells passaged with this frequency as needed.

*Study Approval*

All animal studies were approved by the Stanford University Administrative Panel on Laboratory Animal Care (protocol 27279) and conform to the Guide for the Care and Use of Laboratory Animals published by the US National Institutes of Health (NIH Publication No. 85-23, revised 1996).

*SI References*

1. Jin, S. et al. Inference and analysis of cell-cell communication using CellChat. *Nat. Commun.* **12**, 1088 (2021).
2. Valster, A. et al. Cell migration and invasion assays. Methods 37, 208-215 (2005).
3. Suarez-Arnedo A, Torres Figueroa F, Clavijo C, Arbeláez P, Cruz JC, Muñoz-Camargo C. An image J plugin for the high throughput image analysis of in vitro scratch wound healing assays. PLoS One. 2020 Jul 28;15(7):e0232565. doi: 10.1371/journal.pone.0232565.
